# Adaptive wildlife disease surveillance under uncertainty: field evaluation of a search-theoretic framework for early detection

**DOI:** 10.64898/2026.08.31.748179

**Authors:** Andrew J Bengsen, Sebastien Comte, Lee Parker, Chris Brausch, Fabiola Silva, David M Forsyth, Steven R McLeod

## Abstract

Early detection is critical for successful management of transboundary disease incursions in wild animal populations. However, decisions about how best to allocate wildlife disease surveillance effort must be made under high uncertainty.

We developed and field-tested an adaptive surveillance framework for detecting transboundary disease incursions in wild ungulates in New South Wales, Australia. Key principles guiding its development included accommodating uncertainty, regularly updating search priorities, and a flexible structure that allows the system to be adapted over time. A coarse statewide risk map served as a weak prior describing expected variability in disease incursion risk, loosely focused on foot and mouth disease virus (FMDv). Search values derived from the map were updated every three months based on realised surveillance effort over the preceding 12 months.

Field operations simulating FMDv surveillance collected samples from 1,964 wild pigs (*Sus scrofa*) during two years of development and testing. Most effort was directed to areas of high search value spread across the state. Modelled surveillance system sensitivity ranged from 0.86 to 0.95 over five successive updating cycles. Although the surveillance program was based on FMDv incursion risk, it also fulfilled its secondary goal of detecting unexpected events, including detecting Japanese encephalitis virus in wild pigs before detections in humans and domestic animals.

By combining risk-based surveillance with adaptive updating of search priorities in a modular structure, the framework provided a flexible and generalisable approach for early detection of transboundary animal disease incursions in wildlife populations under high uncertainty.

## Introduction

The successful containment of transboundary and emerging animal diseases is heavily dependent on how quickly they are detected after introduction (Artois et al., 2009; East et al., 2016). However, the earliest stages of an incursion or outbreak are often the most difficult to detect, particularly when they occur in free-ranging wildlife populations (Miller et al., 2022). Delays in detection allow outbreaks to expand, increasing the cost and difficulty of control while also reducing the likelihood of successful containment or eradication (Artois et al., 2009; East et al., 2016). Increasing global travel and trade, environmental change, and changes in the distributions and abundances of wildlife populations are creating new opportunities for transboundary animal diseases (TADs) to spread (Conn and Magalhães, 2024). The development of surveillance systems that can provide reliable early threat detection and situational awareness in wildlife populations has therefore become a critical biosecurity challenge (Stephen and Berezowski, 2025).

Disease surveillance systems that aim to provide early threat detection in wild host populations typically differ in structure, implementation, and interpretation from those that target domestic animals. Wild host populations are often remote, and animals are usually difficult to observe, approach, and sample (Artois et al., 2009). Furthermore, the sizes and distributions of wild host populations are often poorly understood or quantified (Barroso et al., 2024). Consequently, disease surveillance programs for wild hosts are typically more opportunistic, less systematic, and subject to less intensive analysis and reporting than those targeting domestic animals (Artois et al., 2009). These constraints mean that disease surveillance programs for wild animal populations are subject to much greater uncertainty about pathogen occurrence and surveillance effectiveness than programs targeting domestic animals (Nusser et al., 2008).

In Australia, extensive populations of wild pigs (*Sus scrofa*), goats (*Capra hircus*) and six species of deer (Cervidae) create a challenging disease surveillance and management problem. These wild ungulate species are widespread, often occur in remote areas, and can serve as reservoirs for many important pathogens that can harm livestock production, native wildlife, and public health (Bengsen et al., 2014; Cripps et al., 2019; Huaman et al., 2023). Wild pigs are particularly important because they are highly susceptible to several important TADs (Supplementary Figure S1), frequently interact with livestock, and can serve as amplifying hosts (Hone and Pech, 1990; Doran and Laffan, 2005). Establishment of a TAD within a wild ungulate population would greatly complicate disease control and subsequent demonstration of disease freedom. Early detection is therefore likely to be critical for effective containment and a rapid return to pre-incursion conditions.

Notwithstanding the Northern Australian Quarantine Strategy, which aims to provide early warning of biosecurity incursions across the north of the country, the detection of emerging pathogens or diseases in wild ungulates throughout most of Australia has largely relied on passive surveillance (DAFF, 2024). Passive surveillance is an important component of wildlife disease monitoring systems, but its effectiveness depends on a fragile chain of unlikely events, from early detection of visible signs of infection in a wild animal by an educated and motivated observer to the timely follow up by appropriate authorities. Passive surveillance alone cannot be expected to provide the early warning required to enable a rapid and effective management response to a TAD incursion (Artois et al., 2009), especially in a country of approximately 7.7 million km^2^ with a small, spatially concentrated human population (Hone and Pech, 1990). In Australia and elsewhere, these limitations have led to increasing interest in targeted or active surveillance approaches that can reduce the time between an incursion and detection, thereby enabling a timelier response with a greater probability of success (Mörner et al., 2002; East et al., 2013; Miller et al., 2022).

A well-structured active surveillance program can reduce time to detection and provide a quantifiable estimate of surveillance sensitivity (Doherr and Audigé, 2001). However, the question of how surveillance resources should be allocated to best achieve program objectives, given the inescapable high uncertainty characteristic of wild host-pathogen systems, is a recurring challenge. Risk-based approaches can prioritise search effort allocation based on locations where incursions are considered most likely, but they are subject to interacting layers of uncertainty arising from imprecise spatial data, assumptions about pathways of introduction, and imperfect knowledge of host communities and disease dynamics (Nusser et al., 2008; Rodríguez-Prieto et al., 2015). Surveillance systems that focus too narrowly on predicted high risk areas may therefore fail to detect unexpected events. Many studies have proposed spatial risk assessment frameworks to support targeted wildlife disease surveillance (e.g. Tracey, 2010; East et al., 2013; Miller et al., 2022). However, relatively few have reported on their implementation, evaluation, or adaptation. Consequently, there is little evidence available to evaluate how well these approaches can perform in practice under the many constraints and uncertainties characteristic of wildlife disease surveillance.

Search theory can help to guide the efficient allocation of finite surveillance effort by combining uncertainty, imperfect detection, and learning from previous effort to create a dynamic map of likely future search values and priorities. Search-theoretic frameworks combine information about the prior probability of a target’s location, the spatial distribution of realised search effort, and the probability of detection conditional on effort to improve search efficiency and effectiveness (Koopman, 1946; O’Kelly, 2023). Importantly, non-detections provide useful information by reducing the probability that a target remains in a searched area, thereby allowing that probability to be redistributed to unsearched areas and future search priorities to be updated accordingly. Originally developed to locate military assets under extreme uncertainty, search-theoretic approaches are particularly well suited to surveillance problems in which information accumulates gradually through a sequence of imperfect searches. These concepts are directly relevant to surveillance for TAD incursions in wild animal populations, where spatial probabilities of pathogen incursions are uncertain, surveillance sensitivity is imperfect, and resources are insufficient to search all areas intensively.

Here, we report on the development and field evaluation of an adaptive risk-based surveillance framework grounded in search theory, and assess whether a surveillance system that deliberately spreads effort beyond predicted high-risk areas can maintain high system sensitivity while retaining the power to detect unexpected events. The framework was developed from three years of operational experience establishing a TAD surveillance program for wild ungulates in New South Wales (NSW), Australia. Key design principles included explicitly accounting for uncertainty, using search theory to estimate and update the expected information value of surveillance effort through time, balancing expected incursion risk against the need for broad spatial coverage, and maintaining flexibility to respond to changing conditions and new information. The framework is intended not only to detect anticipated threats but also to improve the likelihood of detecting unexpected disease incursions or epidemiological events under high uncertainty.

## Methods

The state of NSW encompasses > 801,000 km^2^ of south-eastern Australia, including 18 bioregions (DCCEEW, 2016). It is Australia’s most populous state, but approximately 85% of the human population resides in coastal cities and towns. In contrast, wild ungulates are widespread across most of the state (Crittle and Millynn, 2023). The need for early TAD detection is particularly acute in NSW because it is the greatest importer of agricultural commodities in Australia (Department of Foreign Affairs and Trade, 2023) and receives the greatest number of international visitors (Australian Trade and Investment Commission, 2026), exposing it to a high risk of TAD-containing material being inadvertently or illegally imported.

Development of the surveillance program commenced in 2022 in response to a perceived increase in the likelihood of a foot and mouth disease virus (FMDv) or African swine fever virus (ASFv) incursion, both of which were expanding into neighbouring countries and trading partners (Robinson, 2022). Although the heightened risk posed by these pathogens provided the initial motivation for the program, the broader intent was to develop a generalisable system that could be adapted to detect any TAD that might become important in future. Given the large number of potential pathogens of concern, the diversity of possible incursion pathways, and the challenges of detecting and sampling wild animals across a vast and sparsely populated landscape, the need to account for uncertainty became a central principle of the program’s design. A second guiding principle was flexibility. The program was designed to be both adaptive, allowing surveillance effort to be reallocated in response to new information, and adaptable, enabling its structure and processes to be easily updated in response to operational experience and changing circumstances. These principles were intended to ensure that the system remained effective, efficient, and sustainable over time.

### Program structure

The initial process for allocating surveillance effort was based on an assessment of spatial variability in the risk of a TAD incursion in a wild ungulate population and estimation of sample sizes required to achieve a high level of confidence in detecting an incursion. Surveillance operations commenced in December 2022, with the aims of: (1) assessing the feasibility and efficiency of different sample collection methods, and (2) developing and adapting procedures through stress-testing across a wide range of geographic and operational conditions. The process for allocating surveillance effort evolved throughout the project in response to information gained through operational experience and deeper consideration of search and decision theory, resulting in a modular structure based on an initial risk map and a repeating search optimisation cycle. We first estimated where an incursion was most likely, then prioritised surveillance in high-value locations, measured how thoroughly those locations had been searched, reduced the future priority of heavily sampled areas, and repeated the process every three months (Figure 1).

**Figure 1:**
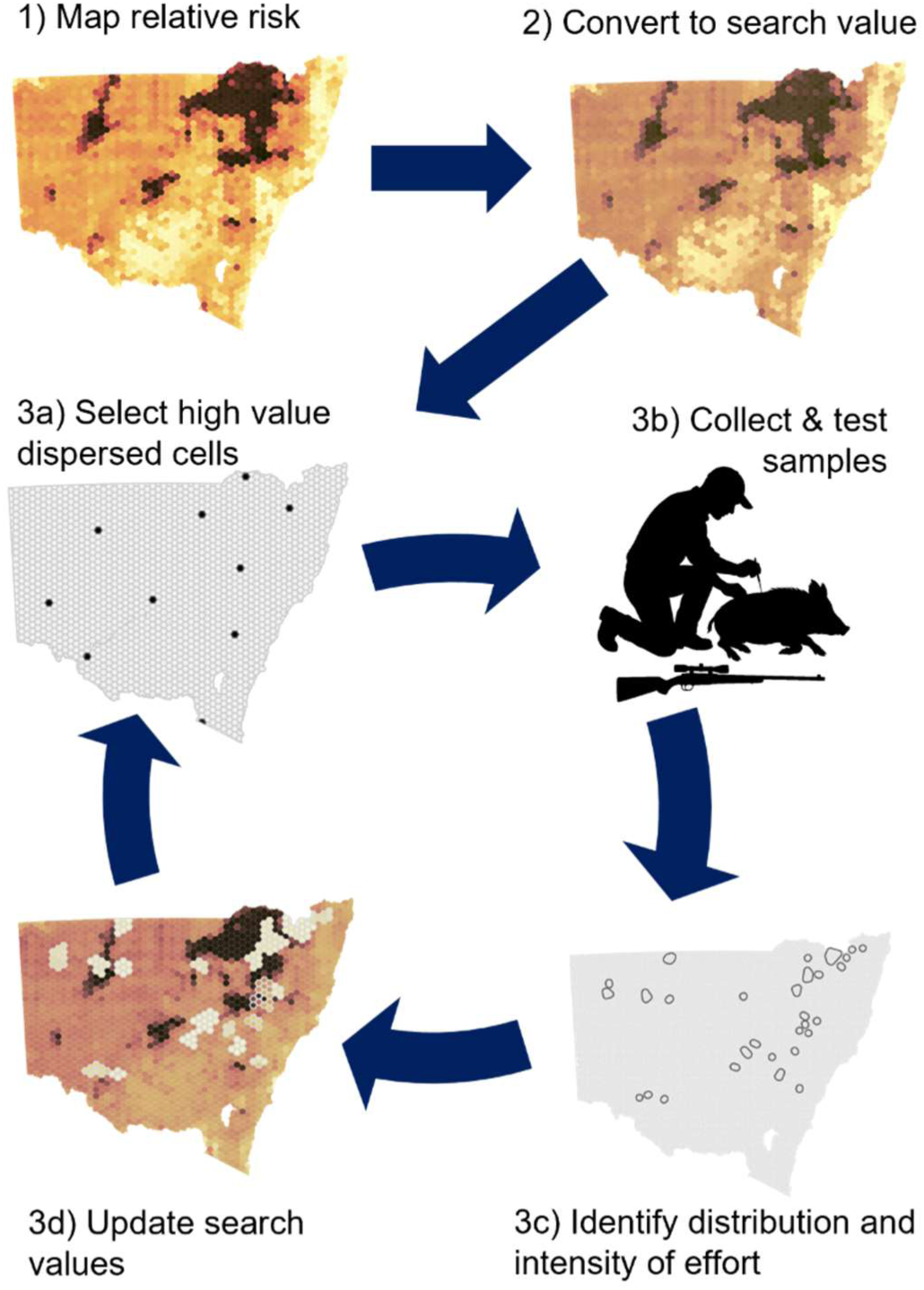
Overview of the adaptive process used to convert prior expectations of relative disease incursion risk (1-2) into a dynamic, actionable set of sites for targeted surveillance. The search optimisation cycle (3a-d) describes the process of updating search targets based on recent effort.

### Risk assessment

The spatial prioritisation of surveillance activities was based on the expected spatial variability in the risk of a TAD incursion in a wild ungulate population and the expected sampling intensity required to achieve a desired level of confidence that an incursion would be detected if it occurred. A surveillance priority map was developed, based loosely on the expected risk of FMDv introduction into a wild ungulate population. FMDv was chosen as a focal pathogen for three reasons. First, it affects multiple species but does not rely on complex transmission pathways within or between species. Consequently, a sampling scheme loosely focused on an FMDv incursion should be more generalisable and provide greater opportunities to detect unexpected events than a scheme tightly focused on a pathogen with more specialised transmission pathways. Second, FMDv occupied a central position in a network of important pathogens and terrestrial livestock host species. This suggested that surveillance strategies designed around FMDv would capture many of the host species, transmission pathways, and wildlife–livestock interfaces relevant to other pathogens of concern, making it a useful surrogate for developing a broadly applicable surveillance system (Supplementary Figure S1). Finally, FMDv has consistently been considered the greatest threat to Australian livestock industries (Hone and Pech, 1990; East et al., 2013) and was a key pathogen of concern at the time due its recent emergence in neighbouring countries and the expected catastrophic consequences of an incursion in Australia (Productivity Commission, 2002; Buetre et al., 2013).

Given the high degree of uncertainty about potential pre- and post-border incursion pathways (Hone and Pech, 1990; East et al., 2013), and the program’s secondary objective of detecting the unexpected, we sought to avoid a tightly focused or over-fitted spatial risk model. Development of the surveillance priority map was, therefore, deliberately uncomplicated. We divided the state into a hexagonal grid comprising 1873 cells, each with an area of approximately 458 km^2^. Each cell was assigned a risk value based on the additive combination of six spatial layers representing different components of the risk of FMDv being introduced into a wild pig population and becoming established (Table 1). Each layer was scaled between 0 and 1, and its contribution to the total risk value was weighted according to its perceived importance as an indicator of the risk of FMDv establishment in a feral pig population.

**Table 1:** Spatial layers and their weightings (summing to 1), used to construct a map of the expected risk of a foot and mouth disease virus incursion in a wild pig population in New South Wales.

| Item | Source format | Weight |
|---|---|---|
| Expected relative wild pig density | Raster | 0.51 |
| Expected relative wild goat density | Raster | 0.08 |
| Expected relative wild deer density | Raster | 0.08 |
| Expected cattle density | Raster | 0.10 |
| Expected sheep density | Raster | 0.08 |
| Location of saleyards, feedlots, and piggeries | Point vector | 0.15 |

Following Hone and Pech (1990) we specified wild pig density as the most reliable predictor of the risk of FMDv incursion and establishment in a wild ungulate population. Wild pigs are recognised as a high-risk host and ideal indicator species for many important pathogens because of their wide distribution, their high susceptibility to pathogen exposure, their ability to produce antibodies, their propensity to mix with livestock, and their role as a major amplifying host that can replicate and shed large quantities of virus (Hone and Pech, 1990; Doran and Laffan, 2005; Barroso et al., 2024). The relative density of wild pigs across NSW in 2023 was mapped on a 5 × 5 km grid using information collected during focus groups with land and resource managers (Crittle and Millynn, 2023). The resulting ordinal data were assigned a numeric rating ranging from 0 to 1, assuming equal intervals between levels. These were averaged across all pixels in each risk map cell to provide a single wild pig density rating scaled between 0 and 1 for each cell. The same process was used to score the expected relative density of wild goats and deer, which could serve as maintenance hosts for FMDv.

Expected densities of cattle and sheep were used to represent potential spillover sources for wild pig populations. Cattle were treated as the main reservoir species driving transmission, while sheep were treated as a secondary reservoir that could maintain infection, with both species capable of infecting wild pigs. Raster data illustrating the expected density of each species in 2020 were downloaded from the Gridded Livestock of the World database (Food and Agriculture Organization, 2024). We used the dasymetric product, in which the spatial distribution of animals across pixels within census units was estimated using random forest models (Gilbert et al., 2018). Estimated animal densities were then averaged across each risk map cell and scaled in the range 0 to 1.

The locations of saleyards, feedlots and piggeries were used to represent the hazard of spillover from high densities of livestock that may have originated from different locations. Active saleyards were identified from an industry report (Meat and Livestock Australia, 2019) and active feedlots and piggeries were identified from an online database (Farm Transparency Project, 2022) and personal knowledge. Risk assessments commonly treat point hazards such as these with a buffer zone of elevated risk. However, each hazard requires subjective choices about the form of the buffer (e.g. radius or decay form). We also expected substantial variability in spillover risk among different saleyards, feedlots and piggeries, depending on their size, connectivity, and biosecurity infrastructure. Given the high uncertainty in the nature of the risk of each point hazard, and in the most accurate way to represent that risk, we simply calculated the density of known point hazards per grid cell, assuming that cells with more combined saleyards, feedlots and piggeries represented a greater risk than similar cells with fewer or no point hazards. Point hazard density was rescaled between 0 and 1.

The six spatial layers were multiplied by their respective weights (Table 1), summed, and rescaled to provide a total relative risk rating for each cell, ranging from 0 to 1. This provided a spatial representation of our expectations of the risk of FMDv incursion and establishment in a wild pig population. We then conducted a sensitivity analysis to assess the effects of varying the weight of each layer except wild pig distribution between 0.1 and 0.5 while holding other weights constant. For each iteration, we calculated the root mean squared error (RMSE) of the revised total risk value relative to the original value.

### Sampling intensity

We used a two-stage, risk-based sampling design (Stevenson, 2021) to estimate the sampling intensity needed to achieve a 95% surveillance system sensitivity (SSe) to detect FMDv non-structural protein (NSP) antibodies in a wild pig blood sample, conditional on FMDv being present in ≤ 5% of cells over a rolling 12-month window. We weighted the risk in each category relative to very low risk cells, such that high-risk cells represented nine times the risk of very low risk cells, and moderate and low risk cells represented seven and three times the risk of very low risk cells, respectively. We assumed laboratory test sensitivity and specificity of 0.97 for an NSP ELISA (Lee et al., 2004; Brocchi et al., 2006; Seeyo et al., 2024).

Given the highly contagious nature of FMDv and its capacity for rapid within-population transmission once infection is established (Fukai et al., 2022), we specified a within-site design seroprevalence of 0.1. This value defines the minimum within-site prevalence at which the surveillance system is intended to achieve a high probability of detection. Required sample sizes were estimated using the *rsu.sssep.rb2st2rf* function in the epiR package (v 2.0.91; Stevenson and Sergeant, 2022). This provided an estimated requirement of 48 sites with 30 pigs tested per site to achieve a target SSe of 95% over a rolling 12-month window, assuming that ≥ 50% of surveillance effort was directed to high-risk cells.

### Surveillance activities

Wild pigs were sampled by the project team through targeted trapping or shooting operations, or opportunistically from aerial shooting and trapping programs conducted by other state government agencies where those operations coincided with the program’s priorities. Sampling operations simulated FMDv surveillance to stress-test and refine operational procedures, evaluate the feasibility and sustainability of ongoing operations, develop collaborative networks with other agencies and private landholders, and collect systematic data on the distribution and prevalence of important emerging and endemic pathogens.

We sought a sample size of 33 pigs per site, comprising the 30 required to meet the objective of 95% SSe and an additional three for redundancy in case of inconclusive samples. Each pig that was sampled was visually assessed for external abnormalities, including lesions consistent with FMD, before taking blood samples and tonsil swabs to test for antibodies indicating exposure to emerging pathogens (e.g. Japanese encephalitis virus, *Brucella suis*) or other pathogens of interest to land managers or government agencies (primarily pathogenic leptospires and *Coxiella burnetii*) (Bengsen et al., 2026a). Occasionally, samples were collected and submitted by partners in other agencies or private enterprise. Although the system was designed around FMDv serological testing, we did not submit samples for FMDv testing because this service was not available at the time. The visual inspections and submission of samples to be tested for other pathogens served as a simulation that allowed us to achieve our development and evaluation goals while also providing data for situational awareness about emerging and endemic pathogens of concern.

### Spatial allocation of surveillance effort

Based on experience gained during initial surveillance efforts, we developed a two-step approach to provide a transparent and repeatable process for allocating future effort. This involved first converting the relative risk rating for each grid cell to an expected search value (Figure 1.2) and then selecting specific cells that provided a balance between high search value and high spatial coverage of the state (Figure 1.3a). The difference between risk and search value is subtle but important. Risk describes where a disease incursion is believed to be most likely. Search value describes where surveillance effort is expected to be most useful after accounting for both risk and previous effort. Consequently, an area may remain high risk but have low search value if it has already been intensively sampled.

To calculate expected search values, we rescaled the total risk rating for each cell so that all cells summed to 1. This represented a weak prior distribution (*p0*) of the probability of FMDv being present in each cell, given its presence somewhere in the state. Based on our initial sample size calculations, we assumed the surveillance system would provide 95% probability of detecting a pig exposed to FMDv, conditional on FMDv having been present at prevalence ≥ 0.1, in a population from which ≥ 30 wild pigs were sampled. This represented the likelihood of detecting FMDv NSP antibodies in a wild pig blood sample when 30 samples were collected from a site where FMDv was present (*d* = 0.95).

The product of the risk rating prior and the likelihood for each cell (*c*) provided the cell’s initial expected search value (*v_c_* = *p0_c_* × *d*). We used the resulting map of expected search values to select cells that balanced the competing objectives of allocating surveillance effort to the most high-risk areas while obtaining broad coverage of the state. We initially selected all cells in the top quartile of search values as candidates for surveillance (n = 468). We then used an iterative risk-weighted selection algorithm to select 10 cells from the candidate set that provided a balance between risk and spatial dispersion. The candidate cell with the greatest expected search value (0.0017) was added to the set of selected cells *S* to initialise the procedure. For each remaining cell in the candidate set, we calculated a selection score:

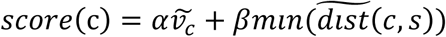

where *v^~^_c_* is the uncentred *z* score for the search value of cell *c, min(dist^~^(c,s))* is the uncentred *z* score of spatial distance to the centroid of the nearest selected cell, and *α* and *β* are weights to balance risk and spatial dispersion, set at 1 and 5, respectively. This weighted-sum approach is commonly used to balance risk and redundancy in multi-objective optimisation problems (Marler and Arora, 2010). The cell with the highest score was then added to *S*, and the loop was repeated until 10 cells had been selected. The resulting set of cells provided a target set that balanced efficient targeting of expected high-risk areas with broad spatial coverage. We selected 10 cells for the set, rather than the full 48 sites indicated by sample size calculations, to minimise spatial clustering. We calculated a new target set of cells every three months based on the distribution and intensity of realised surveillance effort over the preceding 12 months.

### Updating expected search values

In an ideal surveillance program, each sampling operation would target the highest-ranked cell available and then progress to the next highest cell, collecting a full sample size from each cell within the desired time window. In reality, large-scale disease surveillance in wild animal populations is time-consuming, expensive, and requires specialist expertise and access to private properties. Sampling operations must strive for ideal allocation of surveillance effort while balancing constraints set by time, resources, staff fatigue, and site accessibility. This was achieved through a combination of targeted sampling by the surveillance team and selective opportunistic sampling in collaboration with other agencies conducting wild pig control operations in high search value areas.

Selective opportunistic sampling introduces deviations from optimal sampling scenarios that should be accounted for before allocating future surveillance effort. Moreover, search theory dictates that the future search value of any site that has recently been sampled in the search for a non-evasive target should be downgraded, and the search values of remaining sites should be adjusted upward. To achieve this, we developed an iterative procedure adapted from a search method originally developed to search for military assets lost at sea (Richardson and Stone, 1971).

The first step in the search optimisation cycle was to define the spatial distribution of recently searched sites based on spatial clustering (Figure 1.3c). Given that wild pigs are distributed continuously across much of the state, cells on a search value map do not represent distinct populations that can be sampled and analysed independently. Moreover, surveillance operations often crossed adjacent cells. Consequently, a map cell does not represent a distinct site that can be reconciled with the two-stage sampling design. We used a density-based spatial clustering of applications with noise (DBSCAN) algorithm to identify spatial clusters of samples collected during surveillance operations and group them into distinct sites. DBSCAN algorithms are commonly used in machine learning and data mining to define unspecified numbers of arbitrarily-shaped clusters of points, based on a specified neighbourhood radius and minimum number of points to be included in a cluster (Ester et al., 1996; Schubert et al., 2017).

We specified a neighbourhood radius of 10 km and a minimum of 5 sampled pigs to define clusters using DBSCAN, based on samples collected during the preceding 12 months. The 10 km radius was chosen to provide spatial independence given expected activity range sizes and contact rates of wild pigs (Wilson et al., 2023; Proboste et al., 2024). The minimum of 5 sampled pigs provided an expected detection probability estimate of 0.40 within a cluster. The algorithm was implemented using the dbscan package (v 1.2.2, Hahsler and Piekenbrock, 2025) in R (R Core Team, 2024). Each sampled pig was either assigned to a cluster or identified as an outlier if it was spatially isolated or in a group of < 5 pigs. To account for connectivity of sampled pigs with pigs from the broader population, a grid cell was considered to have been sampled if it intersected the 100% minimum convex polygon of a cluster (with an outward buffer of 0.5 × cell width).

The second step in the search optimisation cycle was to estimate the conditional probability of detecting FMDv NSP antibodies in ≥ 1 sample at each site, given the realised sampling intensity within each cluster. The sampling intensity at each cluster was defined as the number of pigs sampled within the cluster during the preceding 12 months. We used the two-stage sampling model to calculate the expected detection probabilities for sample sizes between 1 and 35 pigs (Supplementary Figure S2). These values were then used to downgrade the cell-specific probability of detecting antibodies in a sample from an exposed pig (*d*), from a maximum of 0.97 for a sample of 35 pigs within a site to 0.40 for a sample of 5 pigs. Cells corresponding with clusters that had < 5 samples were considered unsearched, with *d_c_* = 0.

In the third step of the cycle, we adjusted the future search value of all map cells based on their prior search values and sampling effort (Figure 1.3d). Search values were then updated according to:

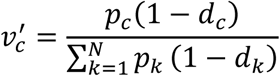

where *v’_c_* is the posterior search value for cell *c*, *p*_c_ is the prior probability that FMDv is present in cell *c*, *d*_c_ is the probability that FMDv would have been detected in cell *c* given the realised sampling effort during the preceding 12 months, and *N* is the total number of cells. The denominator normalises the updated probabilities so that they continue to sum to one across all cells and redistributes probability mass removed from recently searched cells to the remaining cells (Richardson and Stone, 1971). The updated search values (*v’_c_*) were then used to select a new set of high-value cells using the weighted sum algorithm (Figure 1.3a). Cells adjacent to those from which 30 samples have already been collected were excluded from selection. This iterative process ensured that surveillance efforts were continuously re-calibrated, thereby optimising resource allocation and detection probability. This search optimisation cycle was repeated at 3-monthly intervals.

Finally, because no incursions of the target diseases occurred during the study, surveillance performance was evaluated through estimated surveillance sensitivity rather than direct observation of disease detection outcomes. We used the *rsu.sep.rb2st* function from the epiR package (v2.0.91, Stevenson and Sergeant, 2022) to retrospectively estimate SSe for each search optimisation cycle, based on the intensity and distribution of realised sampling effort over the preceding 12 months. Bootstrap confidence intervals for the estimate were estimated by resampling the searched clusters with replacement 2000 times, and recalculating SSe for each iteration. Because these models did not account for the many interacting factors that could influence pathogen detection probabilities in wild host populations (e.g. host density, population connectivity), the resulting estimates should be interpreted as indicators of relative sensitivity, not robust estimates of realised sensitivity.

## Results

### Spatial variability in expected risk

The combined risk map showed high consistency with the underlying wild pig distribution layer (Spearman’s rank correlation coefficient [ρ] = 0.94; Figure 2). When combined with the relative risk categories used to estimate sample size requirements, 49% of cells were rated as high risk (i.e. 9 times greater risk than very low risk), 36% as low risk, and 15% as very low risk. Less than 1% of cells were rated as moderate risk. Sensitivity analysis indicated that substantial changes in layer weights had only minor effects on the resulting risk map (Supplementary Method S1). Increasing the wild goat density layer weight from 0.08 to 0.50 produced the greatest change (RMSE = 0.128), but the spatial pattern of the revised map remained highly consistent with the original (ρ = 0.92). Similarly, increasing the deer layer weight produced an RMSE of 0.121 while maintaining a strong rank correlation (ρ = 0.92).

**Figure 2:**
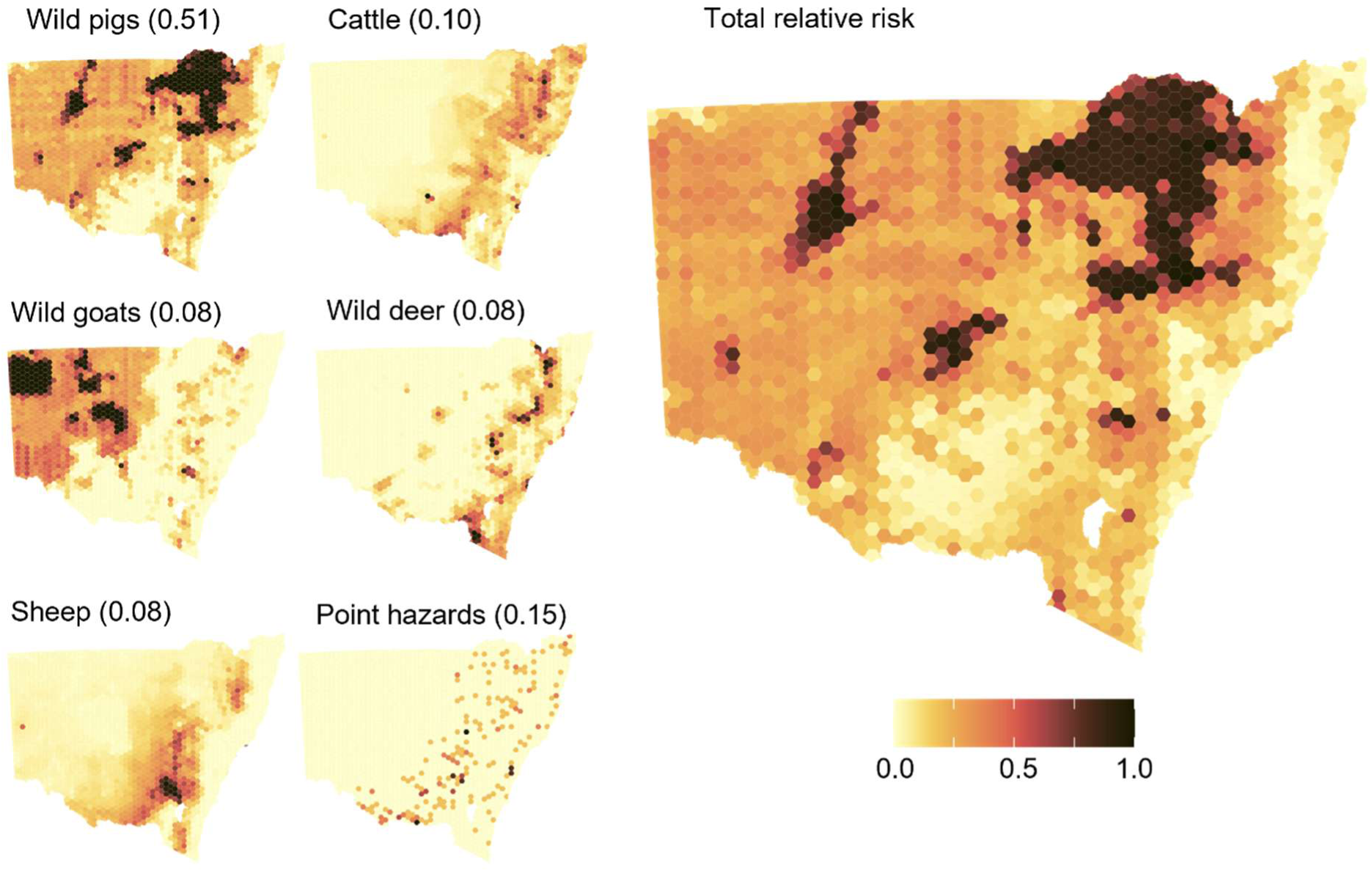
Six spatial layers were combined to produce a map of the expected relative risk of a foot and mouth disease incursion and establishment in a wild pig population in NSW, Australia. Parenthetical figures represent relative risk weights assigned to each layer.

### Operational effectiveness

We ran the first search optimisation cycle in July 2025, based on 945 wild pigs sampled during 42 operations during the preceding 12 months. Most of these samples (52%) were collected during targeted trapping or shooting operations. The remainder were collected opportunistically during population control programs in high search value areas.

The DBSCAN algorithm showed that the 42 operations produced 43 distinct spatial clusters of samples and 16 outlier samples that did not contribute to a cluster. The mean sample size per cluster was 22 pigs, with 26% of clusters achieving a sample size ≥ 30 pigs, corresponding to optimal search intensity. Sample sizes > 33 (maximum = 78) occurred when sites were sampled twice at opposite ends of the 12-month window and when nearby sites were collapsed into a single cluster. Most map cells (74%) that intersected clusters were classified as high risk prior to sampling. The expected search value of most of these cells decreased to very low values after updating, falling below the minimum prior expected search value, although some cells in which far fewer than the target 30 wild pigs were sampled retained low to moderate search values (Figure 3). A further 1,019 pigs were sampled over the next four 3-monthly search optimisation cycles until June 2026, which marked the end of the development and testing phase of the project. These iterations produced similar results to the first, with modelled SSe consistently greater than 0.85, particularly in latter iterations as operational procedures were refined (Table 2). However, only the final iteration achieved the desired modelled SSe of 0.95.

**Figure 3:**
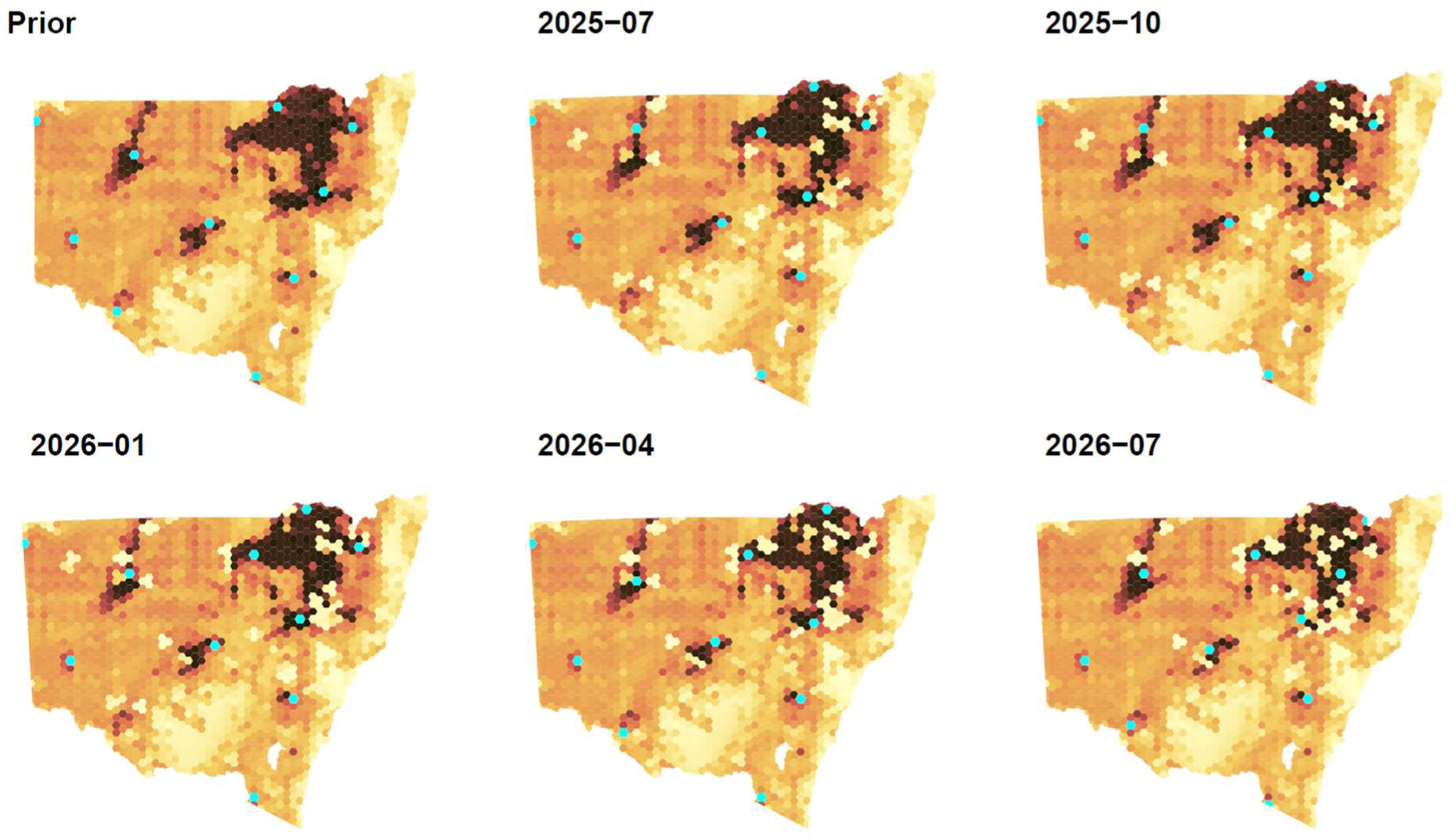
Prior expected search values for each of 1873 grid cells and posterior values adjusted every three months to account for surveillance effort over a rolling 12-month period. Cyan cells indicate locations algorithmically selected to provide a balance of high search value and spatial dispersion.

**Table 2:** Operational results for five 3-monthly iterations of a search optimisation cycle aiming to guide disease surveillance effort based on realised effort over the preceding 12 months. Mean sample size refers to the mean number of samples collected within each cluster, pr ≥ 30 is the proportion of clusters that achieved the target sample size, pr high risk is the proportion of sampled cells that were classified as high risk, and SSe is the model surveillance system sensitivity estimated from realised sampling effort.

| Iteration | Operations | Clusters | Mean n (SD) | pr ≥ 30 | pr high risk | SSe (95% CI) |
|---|---|---|---|---|---|---|
| 2025-07 | 42 | 43 | 22 (15) | 0.26 | 0.74 | 0.90 (0.86, 1.00) |
| 2025-10 | 43 | 36 | 22 (12) | 0.31 | 0.75 | 0.86 (0.82, 1.00) |
| 2026-01 | 56 | 43 | 24 (14) | 0.37 | 0.79 | 0.92 (0.89, 1.00) |
| 2026-04 | 63 | 43 | 24 (14) | 0.42 | 0.86 | 0.93 (0.90, 1.00) |
| 2026-07 | 70 | 48 | 23 (12) | 0.41 | 0.85 | 0.95 (0.93, 1.00) |

### Detection of emerging and endemic pathogens

Blood samples and tonsil swabs collected through the program provided opportunities to detect emerging pathogens. Between December 2024 and February 2025, Japanese encephalitis virus (JEV) RNA was detected in tonsil swabs collected from wild pigs at four widely separated sites. Further detections occurred at two more sites in February 2026.

Serological testing also identified wild pigs exposed to *Brucella suis* across a much broader geographic area than previously recognised, including substantial western and south-western extensions of the pathogen’s known distribution. Detailed analyses of these data and further data on other pathogens are reported elsewhere (Bengsen et al., 2026a).

## Discussion

The adaptive surveillance program evaluated here combines three components of wildlife disease surveillance that are usually treated separately: risk assessment, optimisation of surveillance effort, and regular updating of perceived risk. We integrated these components into a single surveillance framework to address the high uncertainty inherent in many components of a surveillance program. This high inherent uncertainty is one of the greatest recurring challenges in active wildlife disease surveillance.

While many studies have proposed spatial risk assessments as a foundation for efficient and effective targeted surveillance activities (e.g. Tracey, 2010; East et al., 2013; Miller et al., 2022), relatively few have reported on their implementation and evaluation. Our findings contribute to a comparatively small body of work evaluating the practical performance of these approaches in operational wildlife disease surveillance programs (e.g. Desvaux et al., 2021; Schuler et al., 2025).

In a search-theoretic context, risk maps can provide a spatial prior for allocating expected search value. However, spatial risk assessment and mapping can be subject to important biases resulting from data quality issues (e.g. recency, spatial scale, reliability) or crucial information such as unforeseen incursion pathways being missed when the data are compiled (Hone and Pech, 1990; Rodríguez-Prieto et al., 2015; O’Kelly, 2023). Moreover, there are many possible ways in which data layers can be created, processed, and combined.

Consequently, different teams working on similar risk assessment problems can produce very different spatial products (e.g. JEV transmission risk: East et al., 2013; Furlong et al., 2023; Skinner et al., 2025; Walsh et al., 2025). We accepted that we could not reliably predict all important incursion pathways for FMDv or other TADs. To reduce the risk that our prior risk map focused surveillance effort unnecessarily heavily in the wrong areas, we used a deliberately uncomplicated approach to map the expected risk of a FMDv incursion in a wild pig population, informed by few parameters, and mapped at a coarse spatial scale. The resulting risk map was intended to provide a weakly informative spatial prior for assigning expected search values, not a strong prediction of where an incursion would occur. Consistent with this aim, large changes to the weightings of individual risk layers had only minor impacts on the spatial distribution and ranking of risk. This suggests that surveillance priorities were not greatly affected by uncertainty within the spatial model, although models based on different variables could produce substantially different results.

The need to accept uncertainty also implied that surveillance effort should not automatically be concentrated exclusively in areas of highest predicted risk. Concentrating surveillance effort too narrowly could have left the system more vulnerable to unexpected pathogens, unforeseen incursion pathways, or errors in the characterisation of spatial risk. Had we followed an approach focused only on selecting cells with the highest search values, most of our effort would have been concentrated in the north-eastern quadrant of NSW where high wild pig and cattle densities combined to create the highest risk ratings (Fig. 2). Our cell selection process aimed to overcome these risks by balancing expected search value and spatial coverage when allocating surveillance effort. Real-world constraints (see below) meant that it was not possible to sample all selected cells within a candidate set. Instead, we sought to sample as close as possible to as many as possible with targeted sampling operations, while taking advantage of more efficient sampling opportunities in other high search value areas. The spatial clustering of high search value cells across the state meant that when we did sample candidate cells or their immediate neighbours, the next set of candidate cells did not represent a major departure from the spatial pattern of the previous set (Figure 3).

As well as providing opportunities to detect unexpected events, the allocation of surveillance effort across the state should reduce redundancy from spatially concentrated search effort. In contrast to many risk-based programs, our aim was not only to search where an incursion was considered most likely, but also to maximise the likelihood of detecting an incursion despite uncertainty about where it might occur. This balance between exploitation of existing knowledge and exploration of alternative possibilities is a fundamental concept in fields as diverse as foraging ecology (Searle and Hobbs, 2005) and business management (Rojas-Córdova et al., 2023), but is not yet a common feature of wildlife disease surveillance.

The updating of search values relative to estimated detection probabilities in response to realised search effort has been a fundamental feature of searches for targets ranging from lost hydrogen bombs to serial killers (Richardson and Stone, 1971; Rossmo, 2014; O’Kelly, 2023). In our program, variability in detection probabilities arose from sampling error and imperfect sample sizes. Incomplete sample sizes (< 30 pigs) occurred due to many reasons, including low pig population densities, unexpected interruptions to trapping operations, high availability of alternative food sources that reduced trapping efficiency, and opportunistic sampling from population control programs providing fewer samples than expected. Most of the constraints relating to targeted trapping operations conducted by the project team could be attributed to the need to balance the allocation of effort within and among different sampling locations in order to maximise SSe. This is analogous to patch departure decisions in foraging theory, in which a forager aims to allocate search time among disjointed resource patches to maximise food intake and nutrition (Searle and Hobbs, 2005). Refinement of field procedures throughout the development phase of the project resulted in an increase in the proportion of sites achieving target sample sizes without compromising the number of sites, which in turn contributed to increasing SSe (Table 2). Realised sample sizes provided higher SSe than expected because we consistently achieved greater sampling effort from high-risk cells than the 50% conservatively assumed during initial estimation of sample size requirements.

Although the framework was initially developed in response to concerns about FMDv incursions in wild ungulates in NSW, most components of the system are applicable to other pathogens or regions. The processes of risk assessment, estimation of search value, allocation of surveillance effort, and iterative updating are separated into distinct modules, each of which can be modified independently without requiring redesign of the entire system. Consequently, different risk layers, host species, surveillance objectives, or search algorithms could be substituted as needed while retaining the overall decision framework. For example, the same approach could be adapted to other pathogens simply by using a different prior risk map and revising the detection assumptions and test sensitivities. Similarly, existing modules could be replaced or new ones inserted into the existing structure for specific scenarios and objectives. A program aiming to understand the distribution of an emerging disease, for example, could replace the existing search value updating module with one that optimises for uncertainty reduction instead of detection probability. A program aiming to estimate spatially explicit disease freedom probabilities could relax the assumption that disease is present somewhere within the state by updating each cell’s search value independently, rather than redistributing probability among cells.

The discovery of unexpected disease exposure patterns suggests that the program was able to fulfil its secondary objective of detecting unexpected events. Although the program was initially designed around the risk of an FMDv incursion, it successfully detected active JEV infection in wild pigs before detections in humans and domestic animals, despite JEV not being explicitly represented in the spatial risk model. Interestingly, the locations at which JEV was detected were only rated as low to moderate risk in previous JEV spatial risk models (East et al., 2013; Furlong et al., 2023; Skinner et al., 2025; Walsh et al., 2025). The detection of JEV, despite it not being an explicit target of the risk model, supports the conclusion that maintaining broad spatial coverage can create opportunities to detect unexpected events. Similarly, the same surveillance effort generated the largest systematic dataset yet assembled on exposure of wild pigs to *Brucella suis* in NSW. These data have provided valuable situational awareness by revealing substantial extensions of the emerging pathogen’s apparent distribution, identifying previously unrecognised high risk areas, and increasing confidence that large parts of the state remain free from *B. suis* (Bengsen et al., 2026a). These outcomes show that the framework has provided a structured and efficient method for searching broadly and detecting events that were not anticipated during program design.

## Limitations, risks, and future refinements

The surveillance framework presented here represents an important step in delivering protection against TAD incursions in wild ungulate populations in NSW and offers a flexible structure for designing programs with similar goals in other regions. However, there are several limitations that should be considered when interpreting the results of this study.

First, while the surveillance framework and operations performed well during development and field simulation of FMDv surveillance, the program’s effectiveness for detecting a newly introduced pathogen remains untested without a real incursion. Estimates of SSe rely on assumptions about prevalence, host distribution, and diagnostic performance rather than empirical observations from a real outbreak. Importantly, without serological testing, our current ability to detect FMDv relies on visual inspection followed by confirmatory testing, which will have a lower sensitivity that may not be reliably estimable.

Second, detection probabilities were estimated from sample sizes and diagnostic test characteristics using well-established models but did not explicitly account for other drivers of pathogen detectability such as host density or connectivity. A more comprehensive and informative approach would be to simulate pathogen incursion scenarios and surveillance activities within a comprehensive disease transmission and management environment, such as the Australian Animal Disease Spread model (Bradhurst et al., 2015; Bradhurst et al., 2025). This is a major undertaking that was beyond the scope of the present study.

Finally, the framework is intended primarily for early detection and does not currently optimise sampling for delimiting the extent of an incursion, or estimating prevalence, transmission dynamics, or disease freedom following detection. Future work will aim to develop a trigger-response module to estimate an incursion’s extent, but other objectives may be better served by different surveillance designs. In its present form, the framework is likely to be most useful where surveillance aims to provide early threat detection under high uncertainty. Simpler approaches may be preferable where risks are well understood, surveillance resources are abundant for the scale of operations required, or the primary objective is not early warning.

Future refinements to this framework will consider the problems of optimising effort allocation within and among sites and improving sample collection efficiency. This is necessary to consistently achieve the desired SSe. Trapping, in particular, is labour and time intensive (Choquenot et al., 1996). In this program, trapping operations typically required up to two weeks for scouting, free-feeding, and sample collection. During this time, traps are prone to disturbance by people, and heavy rain can make sites inaccessible for days or weeks. A two-week commitment to a single site also represents a considerable opportunity cost. A separate multi-objective optimisation study using our operational data indicated that greater use of small-scale aerial shooting for targeted sampling should improve efficiency, spatial coverage, and program sensitivity (unpublished data).

## Conclusion

Uncertainty is a fundamental and inescapable characteristic of wildlife disease surveillance programs. We developed and trialled a surveillance framework that aimed to improve our ability to detect both expected and unexpected threats, and to remain effective when assumptions are incomplete or inaccurate. The early detection of JEV in wild pigs, and the discovery of previously unrecognised spatial patterns of *Brucella suis* exposure provided strong support for this approach. The modular structure of the surveillance framework provides generalisability and supports sustainability by allowing it to be adapted in response to changing threats, new information, or different management contexts while maintaining a transparent and repeatable process for allocating surveillance effort. More broadly, evaluation of on-ground operations that informed and were informed by the framework showed that a structured process for incorporating imperfect information, learning from surveillance outcomes, and adapting search effort can provide an informative and robust and effective wildlife disease surveillance program in a highly uncertain environment.

## Acknowledgments

Targeted sampling was conducted under approval from DPIRD’s Orange Animal Ethics Committee (OAEC - 0571). Opportunistic sampling from wild pig control programs did not require an Animal Ethics Authority. All pathogen testing was conducted by NSW DPIRD’s Animal and Plant Health Laboratories. Damian Collins (NSW DPIRD Agriculture & Biosecurity) assisted with initial sample size estimation and debugging functions in the epiR package. We gratefully acknowledge the assistance of biosecurity teams at all Land Services Regions and the North Coast Branch of NSW National Parks and Wildlife Service who facilitated sample collection or provided samples from wild pig control programs. We thank Peter Caley (CSIRO), Rhys Powell and Jannene Geoghan (NSW DPIRD Animal Biosecurity) for their insights and advice that improved an earlier report on which this manuscript was based. Contributions of all authors are consistent with the CRediT standard.

## Funding

This project was funded by the New South Wales Government’s Department of Primary Industries and Regional Development.

## Conflicts of interest

The authors declare no conflicts of interest.

## Data availability

The raw data from this study contain animal health records collected on private properties that can not be made publicly available. A reproducible workflow and a synthetic dataset are available on Zenodo (Bengsen et al., 2026b).

## Notes

### Competing Interest Statement

The authors have declared no competing interest.

### Summary of Updates

Changed the title to better reflect the content. Slightly revised the risk assessment to fix a minor scaling error. Updated results in light of the above. Provided DOI for a reproducible workflow.

