## Supplementary Figure S1 for "Adaptive wildlife disease surveillance under uncertainty: field evaluation of a search-theoretic framework for early detection"

Network graph showing the relationships between potential emergency animal diseases and terrestrial livestock (cattle, sheep, horses, pigs, goats) and wild host (pigs, deer, goats) species. Pathogens are drawn from categories 2 and 3 of the Emergency Animal Disease Response Agreement (Animal Health Australia 2025). Foot and mouth disease (FMD) occupies a central location.

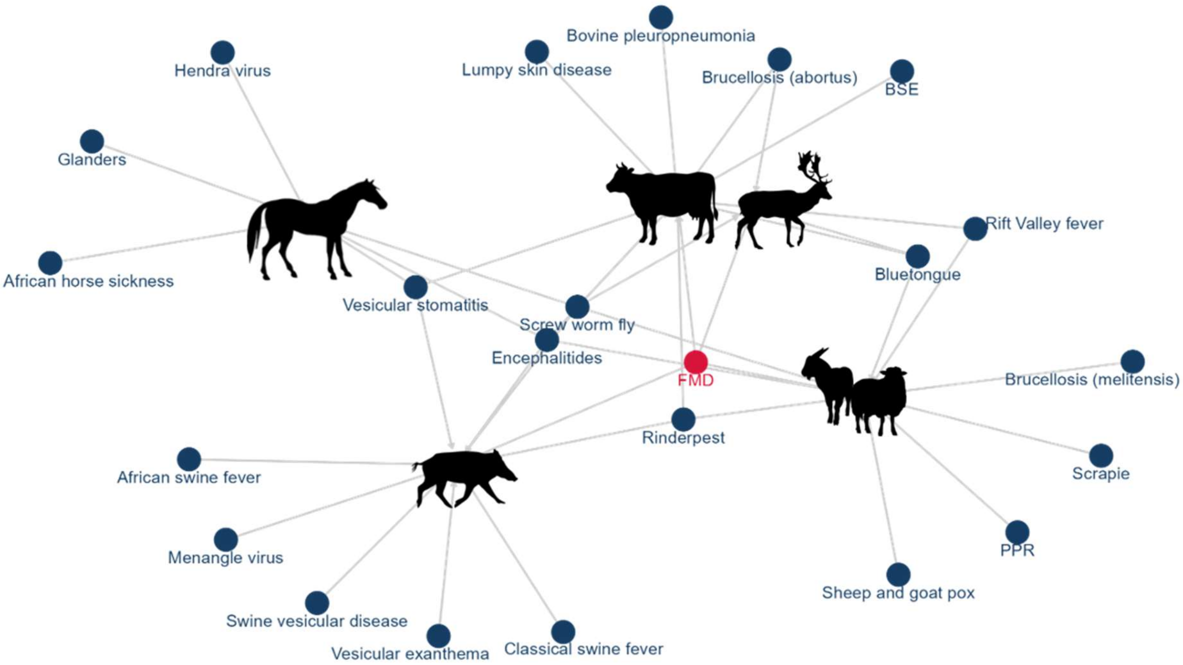


Animal Health Australia (2025) Government and livestock industry cost sharing deed in respect of emergency animal disease responses v25. Animal Health Australia. (Lyneham, ACT).
