## Supplementary Figure S2 for "Adaptive wildlife disease surveillance under uncertainty: field evaluation of a search-theoretic framework for early detection"

Expected within-site FMDv NSP antibody detection probability as a function of sample size, assuming test sensitivity of 0.97 and site seroprevalence of 0.1. 95% confidence intervals were estimated using 10000 bootstrap samples.

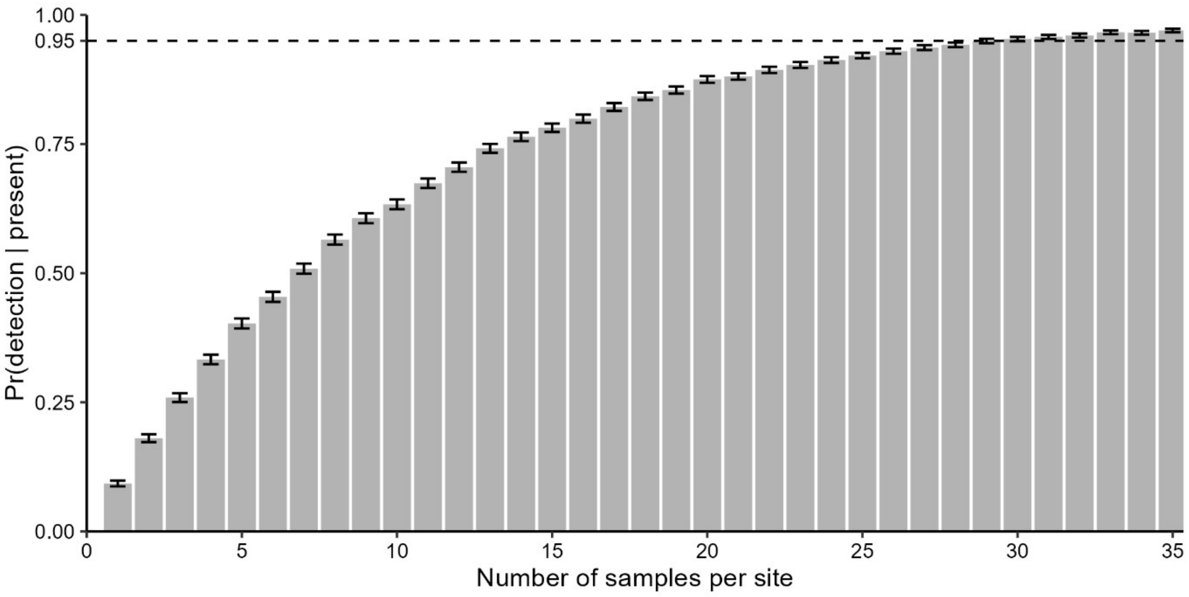
