## Supplementary Method S1 for "Adaptive wildlife disease surveillance under uncertainty: field evaluation of a search-theoretic framework for early detection"

**Supplementary Method S1: Sensitivity analysis of risk layer weightings**

To assess the influence of subjective weighting decisions on the spatial risk map, we conducted a one-at-a-time sensitivity analysis of all risk layers except wild pig density. Wild pig density was treated as the most important risk factor and its weight was held constant at 0.51. For each of the remaining layers (cattle density, sheep density, wild goat density, deer density, and point hazard density), the layer weight was varied from 0.1 to 0.5 in nine evenly spaced increments while all other layer weights were held at their original values.

For each revised weighting scenario, a new total risk map was generated using the same weighted-sum procedure as the primary analysis and rescaled to the range 0-1. The resulting risk surfaces were then compared with the original risk map. Sensitivity was quantified using the root mean squared error (RMSE) between cell-specific risk values in the revised and original maps, and by calculating Spearman's rank correlation coefficient ($\rho$) between the two surfaces. Higher RMSE values indicated greater deviations from the original risk map, whereas high rank correlations indicated that the spatial ordering of cells remained largely unchanged.

This analysis was intended to evaluate the robustness of spatial risk predictions to uncertainty in layer weighting. Because weights were varied independently and were not constrained to sum to one, the analysis should be interpreted as a test of the relative influence of each layer on the spatial distribution of risk.

Increasing the weights of the cattle, goat and deer layers produced the greatest deviations from the original baseline risk map, whereas the sheep and point hazard layers had relatively little effect among the range of weights considered (Supplementary Figure S3). Increasing the cattle density layer weight from 0.1 to 0.5 produced the single greatest deviation from the baseline map (RMSE = 0.049), but the spatial pattern of the revised map remained highly consistent with the original ($\rho$ = 0.96). Increasing the goat layer weight to 0.5 produced a slightly lower RMSE (0.045) but a slightly weaker spatial correlation with the baseline map ($\rho$ = 0.91). Increasing the weights of goat and deer layers tended to produce a more even risk surface than the baseline map (Supplementary Figure S3). The goat and deer layers may have caused greater spatial changes than the cattle layer, despite their lower RMSE, because they were produced from ordinal surfaces with only four possible values whereas the cattle layer was produced from a continuous surface. Increasing the weight of the cattle layer could, therefore, produce large changes in the risk score, leading to larger RMSE, while preserving the risk rankings.

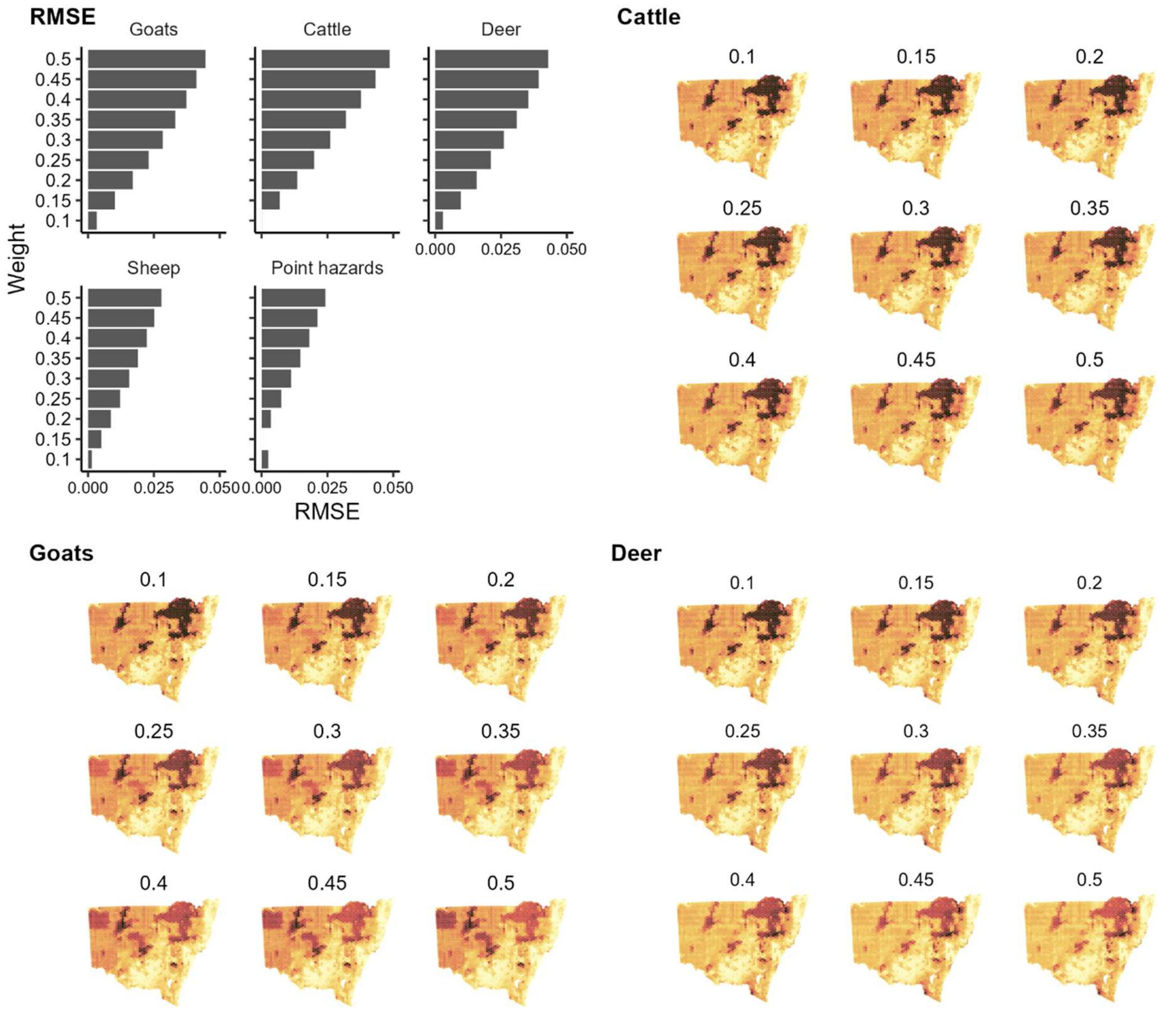


**Supplementary Figure S3: Root mean-squared error (RMSE) describing the magnitude of differences between maps of expected relative disease incursion risk resulting from changing the weightings of five different spatial layers and the baseline risk map, and total risk maps created by varying the weighting of cattle, goat and sheep density layers between 0.1 and 0.5. The weighting for the feral pig density layer was held constant at 0.51.**
